# Hypoxia evokes a sequence of raphe-pontomedullary network operations for inspiratory drive amplification and gasping

**DOI:** 10.1101/2023.11.07.566027

**Authors:** Sarah C. Nuding, Lauren S. Segers, Kimberly E. Iceman, Russell O’Connor, Jay B. Dean, Pierina A. Valarezo, Dale Shuman, Irene C. Solomon, Donald C. Bolser, Kendall F. Morris, Bruce G. Lindsey

**Author notes:** Address correspondence to: Dr. Donald Bolser, Department of Physiological Sciences College of Veterinary Medicine, University of Florida, Gainesville, Florida 32610-0144 USA.

## Abstract

Hypoxia can trigger a sequence of breathing-related behaviors, from tachypnea to apneusis to apnea and gasping, an autoresuscitative behavior that, via large tidal volumes and altered intrathoracic pressure, can enhance coronary perfusion, carotid blood flow, and sympathetic activity, and thereby coordinate cardiac and respiratory functions. We tested the hypothesis that hypoxia-evoked gasps are amplified through a disinhibitory microcircuit within the inspiratory neuron chain and a distributed efference copy mechanism that generates coordinated gasp-like discharges concurrently in other circuits of the raphe-pontomedullary respiratory network. Data were obtained from 6 decerebrate, vagotomized, neuromuscularly-blocked, and artificially ventilated adult cats. Arterial blood pressure, phrenic nerve activity, end-tidal CO_2_, and other parameters were monitored. Hypoxia was produced by ventilation with a gas mixture of 5% O_2_ in nitrogen (N_2_). Neuron spike trains were recorded at multiple pontomedullary sites simultaneously and evaluated for firing rate modulations and short-time scale correlations indicative of functional connectivity. Experimental perturbations evoked reconfiguration of raphe-pontomedullary circuits during tachypnea, apneusis and augmented bursts, apnea, and gasping. The functional connectivity, altered firing rates, efference copy of gasp drive, and coordinated step increments in blood pressure reported here support a distributed brain stem network model for amplification and broadcasting of inspiratory drive during autoresuscitative gasping that begins with a reduction in inhibition by expiratory neurons and an initial loss of inspiratory drive during hypoxic apnea.

## Introduction

Hypoxia associated with a life-threatening event, such as cardiac arrest (1), a seizure (2), or an acute asthmatic episode (3), can trigger a sequence of breathing-related behaviors. An initial increase in breathing effort and frequency is driven by hypoxic stimulation of peripheral chemoreceptors (4). If this tachypnea fails to resolve the hypoxia, the motor pattern transitions to apneusis, accompanied by augmented bursts (a.k.a., sighs) and respiratory depression (5, 6). An ensuing apnea or a dissolution of the respiratory rhythm may be interrupted by gasping, an autoresuscitative behavior characterized by short intense quasi-periodic decrementing bursts of inspiratory activity (7–9). Gasping is a potential biomarker for survival during cardiac arrest (10, 11). The intense inspiratory efforts can generate large subatmospheric intrapleural pressures that yield large tidal volumes resulting in enhanced coronary perfusion and carotid blood flow to the brain (12–14). Accordingly, we postulate that this interdependence effectively “bootstraps,” or coordinates, peripheral chemoreceptor reflexes, the respiratory rhythm, and enhanced sympathetic outflows.

Carotid chemoreceptors operate through multiple paths in the raphe-pontomedullary respiratory network with its embedded, dynamically configured memory system regulated by serotonin (5-hydroxytryptamine, 5-HT) (4). Enhanced brainstem release of 5-HT follows the onset of severe hypoxia (15, 16), and many studies have implicated brain stem deficiencies in serotonin as a contributing factor in sudden infant death syndrome (SIDS) (17–19). SIDS is associated with evidence of hypoxia and failed autoresuscitation (20–23). The action(s) of 5-HT on 5-HT_2A/2C_ and 5-HT_4_ receptors and/or substance P on neurokinin-1 receptors maintain inspiratory motor output during resting breathing and 5-HT can cause the activity of some medullary pre-Bötzinger complex (pre-Böt) neurons to shift to an intrinsic bursting pattern (24–26). *In vitro* neonatal mouse transverse medullary slice studies suggest that gasping depends upon activation of 5-HT_2A_ receptors (27, 28); however, *in situ* arterially-perfused juvenile rat studies do not show a similar role for 5-HT_2A_ receptors in gasping (29, 30). Onset of gasp initiation and recovery of eupnea, however, have been shown to be delayed in animal models with disrupted serotonergic signaling (31–34). Notably, in neonatal mice, acute *in vivo* triggering of synthetic hM4D_i_ (a modified form of the human M4 muscarinic receptor) inhibitory receptors expressed in serotonergic *Pet1-*neurons not only alters baseline cardiorespiratory behaviors but also promotes impaired gasping and reduced survival during apneas (35).

The brain stem network operations that amplify inspiratory drive during hypoxia and gasping are incompletely understood. Previous studies on neuronal and glial signaling mechanisms for augmented breaths and gasping have primarily focused on the ventral respiratory column (VRC) (9, 36–40). The VRC interacts with raphe and pontine circuits that modulate breathing through multiple routes (25, 41–46). Thus, our first objective was to test the hypothesis that hypoxia-evoked gasps are amplified through a distributed efference copy mechanism. This mechanism would be expected to generate coordinated gasp-synchronous discharges in other circuits of the raphe-pontomedullary respiratory network simultaneously.

The chain of VRC inspiratory neurons that generates inspiratory drive is continuously tuned by recurrent excitation and by feed-forward and recurrent inhibitory circuits that actively constrain inspiratory drive (43, 45–50). 5-HT released by the medullary raphe nuclei has been proposed to have a disinhibitory influence on respiratory drive (51, 52). Thus, our second aim was to test the hypothesis that gasping engages a disinhibitory microcircuit within the inspiratory neuron chain to amplify inspiration, thereby enhancing the distribution of gasp drive to multiple targets in the brain and spinal cord.

Our approach to produce hypoxia and gasping was ventilation with a gas mixture of 5% O_2_ in a balance of nitrogen (N_2_). Brain stem neuron spike trains were recorded at multiple sites simultaneously during the evoked changes in the respiratory motor pattern and evaluated for firing rate modulations and short-time scale correlations indicative of paucisynaptic functional connectivity.

Preliminary accounts of some of the results have been reported (53–55).

## Methods

All experiments were performed according to protocols approved by the University of South Florida’s Institutional Animal Care and Use Committee with strict adherence to all American Association for Accreditation of Laboratory Animal Care International (AAALAC), National Institutes of Health and National Research Council, and USDA guidelines. The data comprising this work were part of studies on raphe-pontomedullary network organization and chemoreceptor reflex circuits. Complementary results on the connectivity and central and/or peripheral chemoreceptor-evoked responses of a subset of neurons described here have been reported in previous studies (44–47, 56).

### Surgical protocols

Data were obtained from 6 adult cats (3.0-4.0 kg; 5 females, 1 male) initially anesthetized with isoflurane mixed with air (3-5%) or with ketamine hydrochloride (5.5 mg/kg im); all were maintained with 0.5 – 3.0% isoflurane until decerebration (57). Atropine (0.5 mg/kg, im) was administered at the onset of the experiment to reduce mucus secretion in the airways. Animals were artificially ventilated through a tracheal cannula using a mechanical ventilator; all six cats were bilaterally vagotomized to remove vagal afferent feedback from pulmonary stretch receptors and to permit comparisons with prior work. Immediately prior to decerebration, an anesthetic assessment was performed (44) and cats were neuromuscularly-blocked with pancuronium bromide (initial bolus 0.1 mg/kg followed by 0.2 mg/kg/h iv).

Arterial blood pressure, end-tidal CO_2_, and tracheal pressure were monitored continuously. Arterial PO_2_, PCO_2_, and pH were measured periodically and arterial blood pressure was maintained with solutions of 6% Dextran 70 in 0.9% sodium chloride or 0.04 to 0.1% dopamine (as needed); sodium bicarbonate solution (8%) was used to correct metabolic acidosis. Diphenhydramine hydrochloride (1.8 mg/kg, iv) was administered to reduce mucus secretion in the airways. Dexamethasone (initial bolus of 2.0 mg/kg followed by 4.5 mg/kg/h, iv) was dispensed to help prevent hypotension and minimize brain swelling.

### Changes in chemical drive: Hypoxia

Transient intervals of hypoxia were produced by ventilation with a gas mixture of 5% O_2_ and 95% N_2_ for 90-905 s, then air was reintroduced into the ventilator. In some cats, during the hypoxic exposure and immediately after, arterial blood pressure support was provided by inflation of an embolectomy catheter placed in the descending aorta. For all but one cat, supplemental 100% O_2_ was added to the ventilation gas after blood pressure support was withdrawn.

### Data acquisition and analysis

Descriptions of most of the other methods used have been recently published (46, 50). Signals from a phrenic nerve were monitored. Efferent phrenic nerve discharge was used to identify the phases of breathing and as an indicator of respiratory drive and stimulus effectiveness.

Spike trains were recorded with 3 extracellular electrode arrays (72 total electrodes) at multiple brain stem sites, including the medullary ventral respiratory column (VRC), pons, and raphe; spike trains from individual neurons were isolated using an interactive spike sorting software package (58). Recording sites in the VRC were located 1.4 mm caudal to 6.9 mm rostral to the obex, 3.0 to 4.7 mm lateral to the midline, and 1.9 to 4.8 mm below the surface of the medulla. Recording sites in the pons were located from the caudal border of the inferior colliculus to 1.0 mm posterior to it, 3.7 to 6.1 mm lateral to the midline, and 1.1 to 5.6 mm below the dorsal surface of the pons. Recording sites in the raphe were located 2.2 to 5.6 mm rostral to the obex, within 0.2 mm on either side of the midline, and 1.0 to 5.0 mm below the dorsal surface. Coordinates of recording sites were mapped into the three-dimensional space of a computer-based brain stem atlas derived from Berman (59). Standard firing rate histograms and phase-normalized respiratory cycle-triggered histograms (CTHs) were calculated using activity recorded during a 30-min control period in air; neurons were classified as respiratory modulated if either of two complementary statistical tests rejected the null hypothesis (*p* < 0.05) as previously described (60–62). We further categorized respiratory modulation based on the CTHs (63). Joint time-frequency representations of selected spike train data were generated using an implementation of the S-transform (64) as described in Nuding et al. (56).

Functional connectivity within and among the brain stem regions studied was detected and evaluated with cross-correlation histograms (46, 56), and correlation feature maps, including directed graphs representing inferred connectivity among simultaneously monitored neurons, were generated (45). Significant features in cross-correlograms were identified with Monte Carlo tests using surrogate spike trains (65) with gamma-distributed interspike intervals; FDR < 0.05. The shape parameter of the gamma distribution was estimated from the data (66). A detectability index (DI), calculated as the maximum amplitude of feature departure from background activity divided by the standard deviation of the correlogram noise, that was greater than 3.0 indicated a significant correlogram feature (67, 68).

## Results

The results describe neuronal firing rate dynamics and correlational signatures of functional connectivity during the various respiratory motor patterns induced by transient ventilation with a hypoxic gas mixture (5% oxygen, balance N_2_). For this investigation, a total of 467 neurons (268 VRC, 90 raphe, and 109 pontine cells) were isolated and evaluated in the 6 vagotomized cats studied. Pairs of neurons (n = 20,521) were assessed for evidence of short time scale functional interaction, with 46% of neurons exhibiting functional correlations with at least one other neuron (Table 1). The characteristics and incidence of hypoxia-evoked changes in respiratory motor patterns as indicated by altered phrenic nerve activity are summarized in Table 1.

**Table 1.** Order of hypoxia-induced changes in motor patterns as indicated by altered phrenic nerve activity; not all motor patterns were expressed in every animal. Functional correlations were indicated by significant features in cross-correlations of spike train pairs; features with a detectability index ≥ 3 were considered to be significant (67,68). For example, in Animal 1, 48 of 116 recorded neurons were functionally correlated with at least one other recorded neuron. *, animal died before recovery from hypoxia.

| Animal | Hypoxia Duration<br>(5% O<sub>2</sub>, 95% N<sub>2</sub>) | Motor Pattern Sequence |  |  |  | Number of neurons functionally correlated with at least one other neuron | Number of neurons recorded | % of recorded neurons with at least one functional correlation |
| --- | --- | --- | --- | --- | --- | --- | --- | --- |
|  |  | Augmentation | Depression | Apnea | Auto-resuscitative efforts |  |  |  |
| 1 | 268 s, 305 s (2 trials) | 1<sup>st</sup> | 2<sup>nd</sup> | 3<sup>rd</sup> | 4<sup>th</sup> | 48 | 116 | 41% |
| 2 | 488 s | 1<sup>st</sup> | 2<sup>nd</sup> | — | — | 41 | 110 | 37% |
| 3 | 277 s | 1<sup>st</sup> | 2<sup>nd</sup> | 3<sup>rd</sup> | 4<sup>th</sup> | 22 | 54 | 41% |
| 4 | 905 s | 2<sup>nd</sup> | 1<sup>st</sup> | 3<sup>rd</sup> | 4<sup>th</sup> | 25 | 53 | 47% |
| 5 | 480 s | 1<sup>st</sup> | 2<sup>nd</sup> | 3<sup>rd</sup> | — | 21 | 42 | 50% |
| 6 | 90 s | 1<sup>st</sup> | 2<sup>nd</sup> | 3<sup>rd</sup> | 4<sup>th</sup> * | 60 | 92 | 65% |
| Total |  |  |  |  |  | 217 | 467 | 46% |

### Sequence of motor patterns during ventilation with a hypoxic gas mixture

The sequential generation of motor patterns typically observed in response to ventilation with the hypoxic gas mixture included periods of hypoxia-evoked augmentation (tachypnea), depression (apneusis and augmented bursts), and apnea and gasping - albeit not all motor patterns were noted in all experiments (Table 1). Figure 1 provides the data obtained from *Animal 1* before, during, and after the hypoxic exposure. Shown are the locations of the neuronal spike trains monitored concurrently at multiple sites in the VRC, raphe, and pons (Fig. 1*A*) along with a heat map of neuronal firing rates measured simultaneously in 109 neurons before, during, and after hypoxia, which highlight the rate modulations during the stimulus period (Fig. 1*B, top*); dark tones indicate minimal firing rates and brighter tones indicate faster firing rates. Firing rate histograms for a subset of the neurons during the same time period, together with integrated phrenic nerve activity, blood pressure, and end-tidal CO_2_, document changes in the drive to breathe. Each of the three brain stem regions included neurons with firing rates tightly coupled to the gasp motor burst pattern evident in the phrenic record; note that bursts from neurons P4, V1, and R3 were synchronous with the bursts in phrenic discharge (Fig. 1*B*, *bottom*). Control respiratory CTHs show that neurons with either inspiratory or expiratory discharge patterns exhibited gasp-associated modulation of firing rate (Fig. 1*C*). In general, during hypoxic exposure, we observed a period of augmented breathing characterized by increased phrenic amplitudes and/or tachypnea, followed by a depression which could include apneusis, then finally apnea and gasping; vertical lines though the firing rate heat map and histograms separate these changes in motor patterns (Fig. 1*B, top*). Not all animals exhibited each phase and the phases were not always in this order; Table 1 details the presence and order of periods of the augmentation, depression, apnea and gasping motor patterns for each animal during hypoxic ventilation.

**Figure 1.**
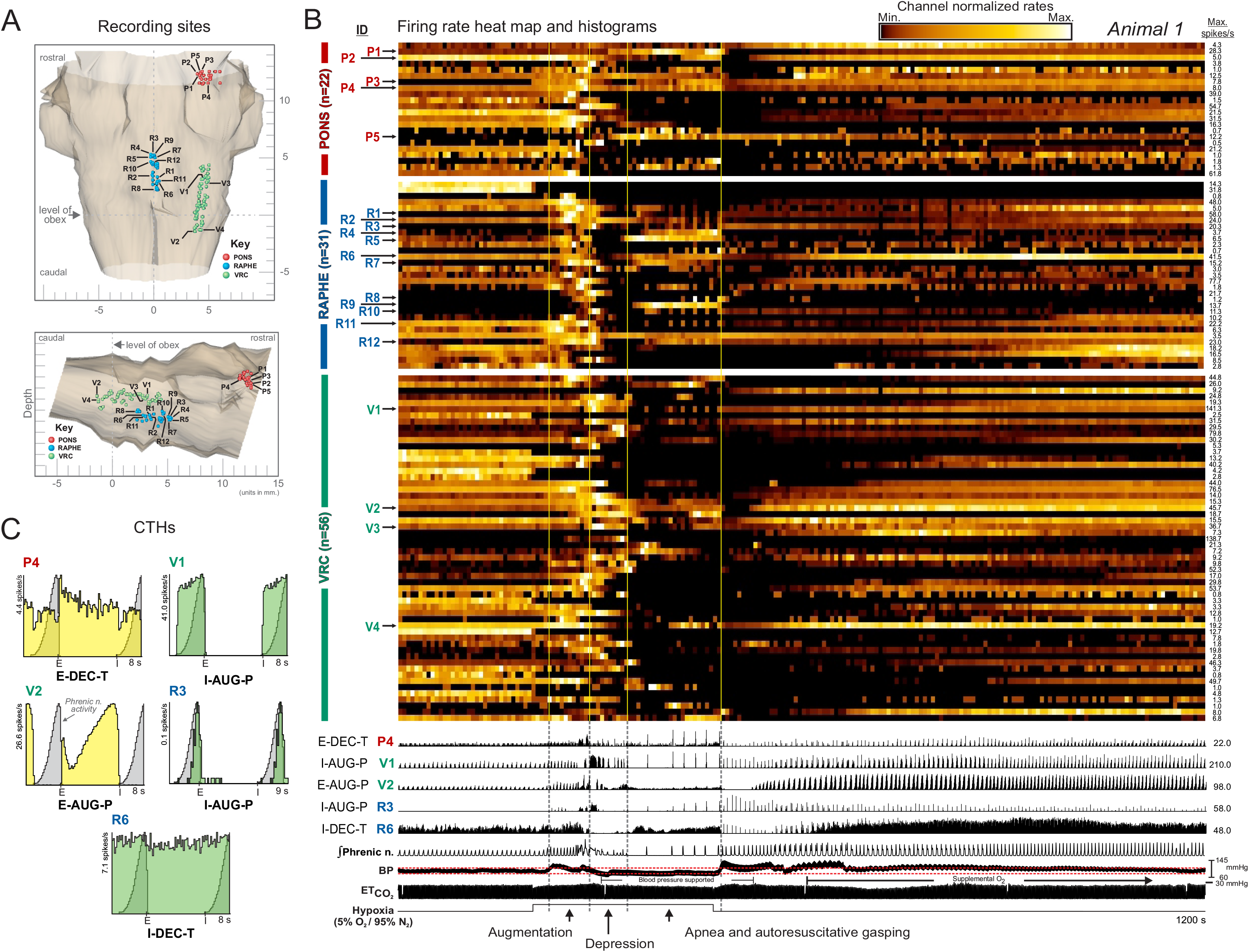
Recording sites and neuron firing rates during a hypoxia-evoked sequence of motor patterns produced by ventilation with a gas mixture of 5% O_2_. *A.* Dorsal (*top*) and sagittal (*bottom*) views of the location of recording sites of neurons in *Animal 1* monitored simultaneously in the VRC (green), pons (red), and midline raphe (blue) mapped into the brain stem atlas. *B.* Firing rate heat map (*top*, with luminance proportional to respective normalized rates) and firing rate histograms for selected neurons, together with integrated phrenic nerve activity, blood pressure, and end-tidal CO_2_ (*bottom*); hypoxic gas ventilation for 268 s. Brain stem regions and neuron labels are to the *left* of the traces. Black bar beneath blood pressure trace denotes period of blood pressure support with inflation of an embolectomy balloon catheter in the descending aorta; red dashed lines in blood pressure trace indicate blood pressure range seen during preceding control interval. Note that 100% O_2_ was added to the ventilation gas after blood pressure support was withdrawn. These data document changes in rate and pattern as the drive to breathe is first augmented and then depressed to an apneic state interrupted by a gasping motor pattern. Vertical lines through the heat map and rate histograms separate periods of respiratory augmentation, depression, and apnea and autoresuscitative gasping motor patterns. *C*. Selected phase-normalized respiratory cycle-triggered histograms (CTHs) show the firing patterns of one pontine cell (P4), 2 VRC neurons (V1, V2), and one midline raphe cell (R6) during eupneic control; cell R3 was not active during control, so data for its CTH was collected during and after the hypoxic exposure. Plot tic marks indicate the onsets of expiration I and inspiration (I).

Higher temporal resolution firing rate histograms for the same animal show gasp-related bursts indicative of widespread engagement of the respiratory network during and immediately following the hypoxic exposure trial (Fig. 2*A*, *arrows*). Gasp-patterned bursts in phrenic motor output were associated with distinct and diverse firing rate profiles in VRC inspiratory neurons. For example, high firing rates in neuron V1 were temporally congruent with transient declines in firing probability in neuron V3 (Fig. 2*A, inset*, and *B)* and *vice-versa* (Fig. 2*C*). Fig. 2*B* shows an augmented phrenic burst (above the dashed line) at the end of the inspiratory activity coincident with increased neuron V1 activity (*asterisk*), yet neuron V3 was quiet during the augmented phrenic burst (*red arrow*). Conversely, at times when neuron V1 has decreased firing (Fig. 2*C*, *left box*), neuron V3 continues to burst without diminution. During the apneustic burst in Fig. 2*C* (right box), however, the neuron firing activities reverse, and neuron V1 increases firing while neuron V3 firing rate declines. The cross-correlogram constructed from activity prior to hypoxia for this pair of neurons had an asymmetrical primary central peak (Fig. 2*D, arrow*) and adjacent secondary bilateral troughs and peaks; the average of the phrenic nerve signal triggered by spikes in neuron V1 also displayed bilateral peaks and troughs (Fig. 2*E*). These high frequency oscillations of firing probability are commonly observed in pairs of inspiratory neurons (HFOs: 60-110 Hz; (69)).

**Figure 2.**
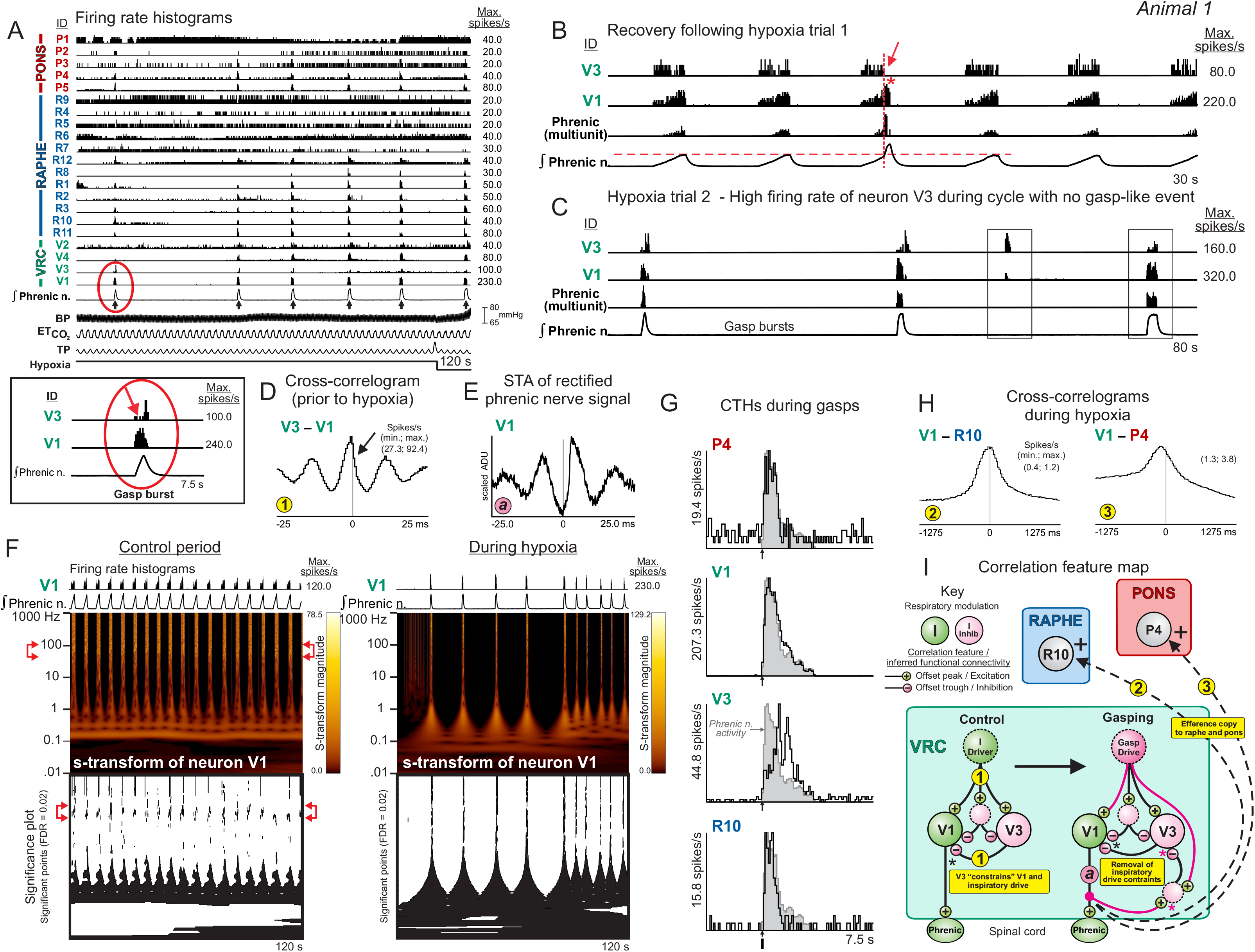
Dynamic neuronal firing patterns during hypoxia-evoked gasping. *A.* Firing rate histograms of selected neurons recorded in *Animal 1* detail transient bursts in the three monitored regions of the brain stem respiratory network during gasp discharges seen in phrenic motor output (*arrows*), consistent with distributed efference copies of inspiratory drive to pontine and medullary raphe neurons. Inset shows detail of reduced activity in neuron V3 (*red arrow*) during peak burst rate in cell V1. *B*. Firing rate histograms during recovery from hypoxia exposure trial 1 show absence of spiking in neuron V3 (*red arrow*) and increased firing in neuron V1 (*red asterisk*) during a period of augmented phrenic burst activity. Also shown is a measure of multiunit phrenic activity derived from the integrated phrenic nerve signal. *C.* Firing rates during gasp bursts during hypoxia exposure trial 2 show reciprocal rate changes in the two neurons during a “missed” phrenic event (*left box*) and during a subsequent apneustic burst (*right box*). *D.* Cross-correlogram for neurons V3 and V1 includes an asymmetrical central peak flanked by trough-peak features; data collected before hypoxic trials. The detectability index (DI) value for the central peak is 19.4. Number of spikes for each neuron used to calculate cross-correlograms: V1: 102,124; V3: 30,059. *E.* Spike triggered average (STA) of full wave rectified left (contralateral) phrenic nerve activity triggered by VRC cell V1. *F.* Firing rate histograms and corresponding s-transform time-frequency representations for neuron V1 during control and gasp motor patterns; s-transform magnitude values range from 0 to 78.5 during control, 0 to 129.2 during hypoxia. *G.* CTH show augmenting pattern for neuron V3 and decrementing patterns for neurons V1, P4, and R10 during gasping. *H.* Cross-correlograms constructed from activity during and after the hypoxic exposure trials are consistent with efference copies of inspiratory drive. The DI values for the peaks in each histogram: *2:* 10.95; *3:* 3.27. Number of spikes for each neuron: P4: 7313; V1: 102,124; R10: 1,259. *I*. Correlation feature map and other hypotheses suggested by the results and prior work; see text. For this and other feature maps, numbers in small yellow circles denote cell pair correlograms (and letters in pink circles denote spike-triggered averages) shown in the current figure.

The s-transform time-frequency representation for the spike train of neuron V1 had a significant magnitude in the HFO band during control inspiratory bursts (Fig. 2*F left, bracketed arrows*); a broader spectrum of significant frequencies was present during gasping bursts (Fig. 2*F right*). During the interval that included hypoxia-evoked gasping, synchronously discharging pontine and raphe neurons P4 and R10, respectively, had decrementing inspiratory profiles similar to VRC neuron V1 (Fig. 2*G*); neuron V3 had an augmenting inspiratory pattern. Cross-correlograms calculated from these data for trigger neuron V1 and targets R10 and P4 had broad central peaks (Fig. 2*H, 2 and 3*). Collectively, these results are consistent with contemporary models of the VRC inspiratory neuron chain and suggest that neurons V1 and V3 received excitation and delayed inhibition via upstream I-driver neurons under control conditions (Fig. 2*I, left*). The results also support the hypothesis that efferent copies of an enhanced inspiratory drive produced by disinhibition were transmitted to the raphe and pons (Fig. 2*I, right and correlograms 2 and 3*).

### Functional connectivity of neurons with hypoxia-evoked tachypneic, apneustic, and augmented bursts, and gasp-like motor patterns

In some experiments, ventilation with the hypoxic gas mixture evoked an apneustic motor pattern with intermittent augmented bursts, but no gasping. Figure 3 provides the data obtained from *Animal 2* to illustrate this hypoxia-evoked motor pattern and the associated changes in neuronal activities. During the initial period of enhanced inspiratory drive, simultaneously monitored pontine and VRC neurons had firing patterns with active intervals or pauses that matched the durations of concurrent apneustic inspiratory bursts (Fig. 3*A, left box*). The firing pattern of I neuron V8 changed from augmenting during the respiratory cycles immediately prior to the hypoxic exposure to decrementing during the apneustic cycles; although the neuron fired throughout apneustic inspiration, its highest rate occurred during the first half of the I phase (Fig. 3*A*, *top red arrow*). In contrast, the firing rates of two decrementing inspiratory neurons, V9 and V10, were more constant throughout the apneustic I phase, and they maintained their overall decrementing pattern of activity (Fig. 3A, *bottom red arrows*). Concurrently, augmenting expiratory (E-Aug) neuron V5, post-inspiratory (or decrementing expiratory, E-Dec) neuron V6, and tonic expiratory (t-E) neuron V7 exhibited reduced firing probability throughout the apneustic inspiratory phase (Fig. 3*A*, *red asterisks*).

**Figure 3.**
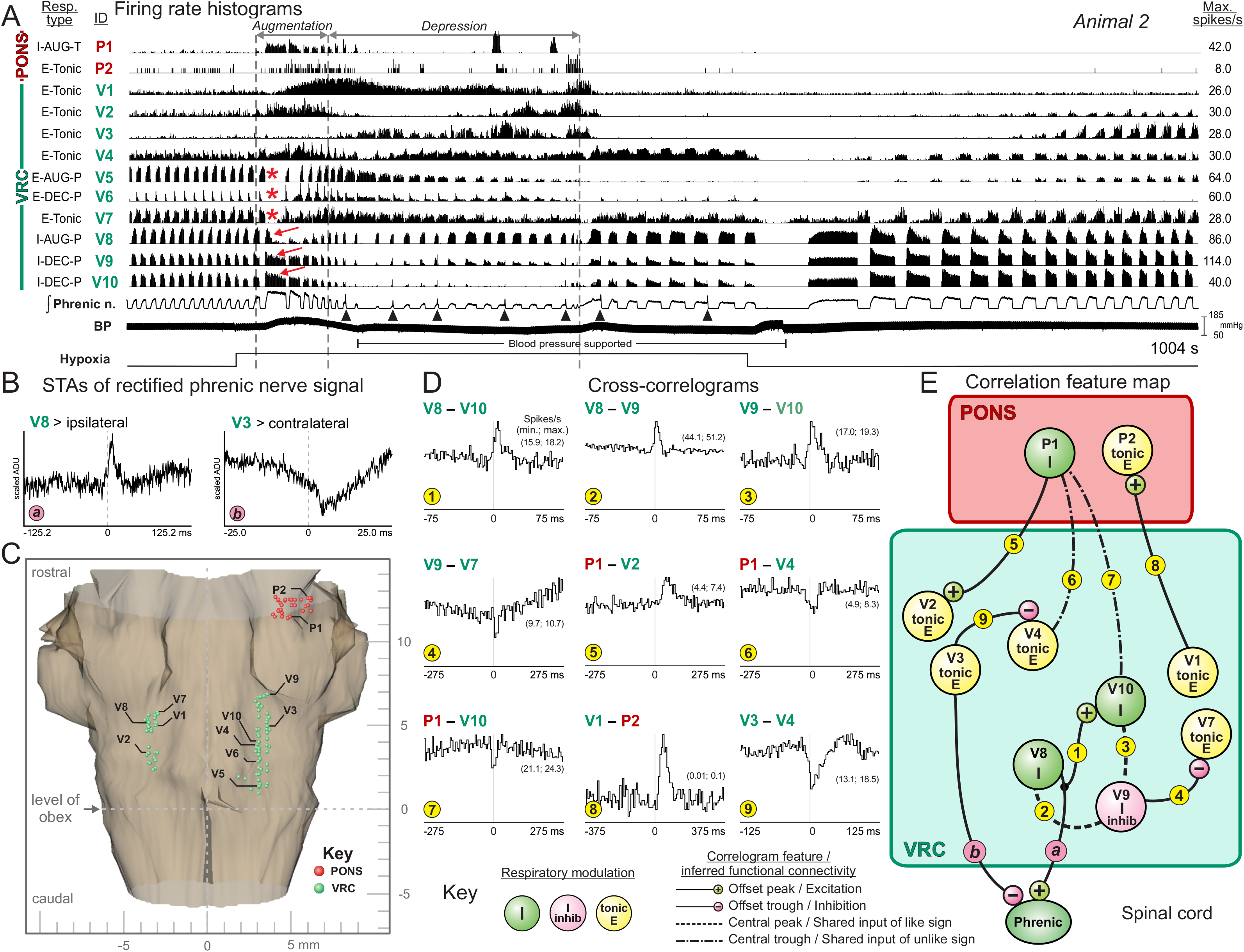
Hypoxia-evoked apneustic motor pattern with augmented bursts. *A*. Firing rate histograms of pontine and bilateral VRC neurons together with integrated phrenic nerve and arterial blood pressure before, during, and after ventilation with 5% O_2_; apneustic patterns and augmented bursts (*arrowheads*) were detected. Black bar beneath blood pressure trace denotes period of blood pressure support with inflation of an embolectomy balloon catheter in the descending aorta. Red arrows and asterisks mark cycles discussed in text. *B*. STA of rectified phrenic nerve activity show offset peak and trough features for trigger neurons V8 and V3, respectively. *C.* Dorsal view of the location of recording sites of neurons in *Animal 2* monitored simultaneously in the VRC (green) and pons (red) mapped into the brain stem atlas. *D*. Cross-correlograms for indicated pairs of neurons. DI values for troughs or peaks in each histogram: *1*: 6.29; *2*: 13.09; *3*: 6.49; *4*: 3.51; *5*: 5.32; *6*: 5.49; *7*: 3.59; *8*: 6.85; *9*: 8.78. Number of spikes for each neuron: P1: 9,824; P2: 384; V1: 51,534; V2: 17,534; V8: 201,037; V7: 109,890; V4: 52,680; V3: 133,080; V10: 68,479; V9: 198,262. *E.* Correlation feature map; see text for details.

As the hypoxic episode continued, phrenic activity was depressed and augmented bursts located near the end of the inspiratory activity occurred intermittently (Fig. 3*A, arrowheads below integrated phrenic nerve signal*). The average of the rectified phrenic nerve signal triggered by I-Aug neuron V8 had an offset peak with a positive lag, a result consistent with a pre-motor function (Fig. 3*B*, a). In contrast, the average of the rectified phrenic nerve signal triggered by t-E neuron V3 had an offset trough suggestive of functional inhibition of inspiratory drive (Fig. 3*B*, b). The firing rates of t-E neurons V3 and V4 fluctuated as the hypoxic exposure continued, with highest firing rates in each tending to occur at different times. Notably, an offset cross-correlogram trough supported an inhibitory action of neuron V3 upon neuron V4 (Fig. 3*D*, 9).

Cross-correlation analysis of other neuron pairs (Fig. 3*D*) revealed features suggestive of various VRC and pontine network operations and supported observed firing rate changes. The functional interactions inferred from correlogram features are summarized in the corresponding correlation feature map (Fig. 3*E*). VRC t-E neuron V1 triggered a cross-correlogram with pontine t-E neuron P2 featuring an offset peak (Fig. 3*D*, 8). The offset peak in the correlogram for pair V8-V10 (Fig. 3*D*, 1) and asymmetric central peak features for pairs V8-V9 and V9-V10 (Fig. 3*D*, 2-3) were consistent with interactions between, or shared influences among, this trio of neurons. We note that neurons V10 and V9 had decrementing discharge patterns during preceding control inspiratory bursts, and that the cross-correlogram triggered by V9 for target t-E neuron V7 had an offset trough feature consistent with a functional inhibition of the t-E neuron that likely continued to operate during the apneustic bursts (Fig. 3*D*, 4) as shown in Fig. 3*A* (red asterisks). Cross-correlogram features for pontine trigger neuron P1—also active during apneustic bursts (Fig. 3*A*)—included an offset peak with target t-E neuron V2 (Fig. 3*D*, 5) and central troughs with targets V4 and V10 (Fig. 3*D*, 6-7). These latter features suggest unobserved shared influences having opposite actions upon the elements of each pair.

### Differential modulation of neuron firing rates during tachypnea, hypoxic gasping, and gasp-associated step increments in blood pressure

Ventilation with hypoxia in *Animal 3* evoked a sequence of motor patterns that began with augmented inspiratory drive and tachypnea followed by apneusis and respiratory depression, leading to apnea and gasping (Fig. 4*A*). The VRC neurons monitored (Fig. 4*B*) during this motor pattern sequence included decrementing and augmenting inspiratory and expiratory neurons (Fig. 4*C, F*). In addition to an increased respiratory cycle frequency, differential changes in the firing rates of the I-Aug neurons were apparent during tachypnea (Fig. 4*C*). The peak rate of neuron V5 increased, whereas that of neuron V4 declined, in some cycles to such an extent that only two small bursts were apparent at the start and end of the inspiratory phase (Fig. 4*C*, inset 2; *asterisks*). Concurrent with these altered patterns, the peak rate of E-Aug neuron V2 declined as that of I-Dec neuron V3 increased (Fig 4C).

**Figure 4.**
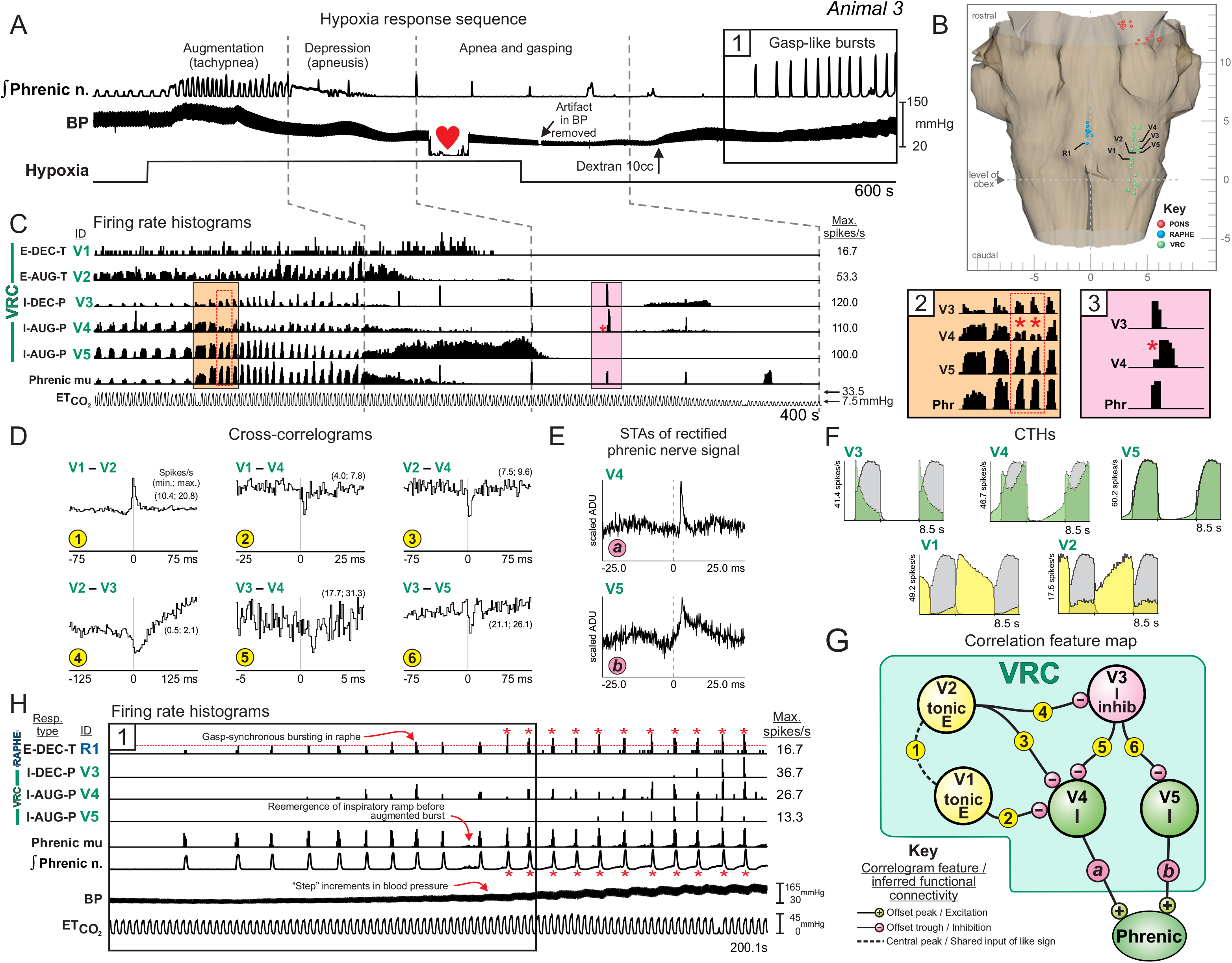
Apneustic, tachypneic, and gasp motor patterns during hypoxia induced by ventilation with 5% O_2_. *A*. Integrated phrenic trace, blood pressure and stimulus marker showing interval before, during, and after hypoxic gas ventilation. Black arrow denotes time of 10cc dextran bolus. Heart symbol indicates time of blood collection for blood gas measurement (the trace is flat because blood was diverted from the pressure transducer during collection): PO_2_ = 31.8 mmHg; pH =7.41; PCO_2_ = 25.6 mmHg. *B*. Dorsal view of the location of recording sites of neurons in *Animal 3* monitored simultaneously in the VRC (green) and raphe (blue) mapped into the brain stem atlas. *C*. Firing rate histograms showing changes in discharge patterns of VRC neurons during hypoxia together with multiunit (mu) phrenic activity and end-tidal CO_2;_ the data segment shown here corresponds to the first 400 s of the segment shown in *A*. Insets 2 and 3 show details of firing dynamics highlighted in *C*. *D*. Cross-correlograms for the indicated pairs of neurons. DI values for troughs or peaks in each histogram: *1*: 25.01; *2*: 7.81; *3*: 6.64; *4*: 7.41; *5*: 6.09; *6*: 4.92. Number of spikes for each neuron: V3: 53,680; V1: 59,942; V4: 113,205; V5: 175,482; V2: 63,600. *E*. STA of rectified ipsilateral phrenic nerve activity triggered by neurons V4 and V5. *F*. Phase-normalized respiratory CTH show the firing patterns of five VRC neurons during eupneic control. *G.* Correlation feature map for indicated neurons; see text for details. *H.* Detail of panel *A* (box 1) on an expanded time scale showing firing rate histograms of raphe and VRC neurons together with multiunit and integrated phrenic nerve activity during gasp-like bursts following the interval of hypoxia exposure; blood pressure and end-tidal CO_2_ are also shown. Note step-wise increments in systemic arterial blood pressure associated with each burst in VRC and raphe neurons (red asterisks).

A central peak feature in the cross-correlogram for t-E neurons V1 and V2 suggests a shared source of drive (Fig. 4*D, 1*); neither neuron was active during the period of apnea and subsequent gasps (Fig. 4*C*). The cross-correlograms triggered by neurons V1 and V2 with target inspiratory neurons V4 and V3 were characterized by offset troughs (Fig. 4*D*, *2-4*), implying convergent as well as divergent functional inhibitory influences. Cross-correlograms triggered by neuron V3 had offset trough features consistent with divergent functional inhibition of neurons V4 and V5 (Fig. 4*D*, *5-6*); the neuronal activities presented in insets 2 and 3 (Fig. 4*C*, *asterisks*) show cycles in which a reduction in the firing rate of neuron V4 coincides with the peak rate of neuron V3. Collectively, these data support the hypothesis that loss of activity in neurons V1 and V2 during apnea resulted in reduced inhibition of neurons V3 and V4 and led to differential tuning of the firing rates of neurons V4 and V5, both of which triggered phrenic signal averages featuring offset peaks suggestive of premotor functions (Fig. 4*E*).

These correlation features are summarized in Fig. 4G. With the return of air ventilation and support of blood pressure, gasp-like bursts reemerged in the phrenic nerve, accompanied by step-like increases in arterial blood pressure (Fig. 4*A*, *box 1*; Fig. 4*H red asterisks*). Concurrent bursts occurred in raphe neuron R1 and VRC neurons V3, V4, and V5; increased firing rates in these neurons were associated with the increases in blood pressure.

A similar sequence of respiratory depression and apnea leading to gasping, with the gasp bursts coordinated with increments in blood pressure was observed in *Animal 4* (Fig. 5). Ventilation with hypoxia evoked an initial apnea (not shown) followed by gasping discharges intermingled with augmented breath sequences; peak firing rates declined with successive bursts in some neurons and increased in others (Fig. 5*A*). Several of the larger gasp-patterned phrenic bursts followed a series of smaller, yet successively incrementing, inspiratory phase discharges (e.g., VRC neuron V12). During the latter part of the hypoxic exposure period, step-like increases in blood pressure developed (Fig. 5*A*) and these were associated with increased gasp-burst amplitude in the phrenic nerve (*red asterisks*) and gasp-synchronous firing in raphe (I neuron R5) and VRC (I neurons V7, V8, V9, V10, V11, and V12) neurons. Slow oscillations in the firing rates of other raphe (R1 and R4) and VRC (e.g., V1, V3, and V4) neurons synchronized with the fluctuations in blood pressure were also detected during this “late” response period (Fig. 5*A*). Some neurons exhibited a greatly reduced firing rate during the late response (neuron V5) or stopped firing altogether (neuron V6). The cross-correlogram for augmenting inspiratory neurons V7 and V12 had a central peak feature consistent with shared excitation (Fig. 5*C*, 1); offset peaks in their STA of the rectified phrenic nerve signal suggest a premotor function for the two neurons (Fig. 5*B*, a, b). The dual peaks in the correlogram for neuron pair V12–V11 (Fig. 5*C*, 3) were consistent with a reciprocal excitatory relationship (Fig. 5*A*), one operation appropriate for a role in inspiratory drive amplification (Fig. 5*D*, 3). The hypothesis that inspiratory chain mediated disinhibition contributes to gasp drive amplification was supported by correlational signatures of functional connectivity among the monitored neurons (Fig. 5*C, D*). For example, the offset trough feature in the cross-correlogram triggered by neuron V6 for target neuron V7 (and the inferred functional inhibition of V7 by V6; Fig. 5*C*, 2) was consistent with the activities of these two neurons during the early period of response when the firing patterns of the 2 neurons are asynchronous (Fig. 5*A*, 2; *red double-headed arrows*), as well as during the late period when the firing rate of inhibitory neuron V6 declines precipitously and the gasp-synchronous firing of neurons V7 becomes greater, indicating amplification of inspiratory drive via disinhibition (Fig. 5*D,* 2).

**Figure 5.**
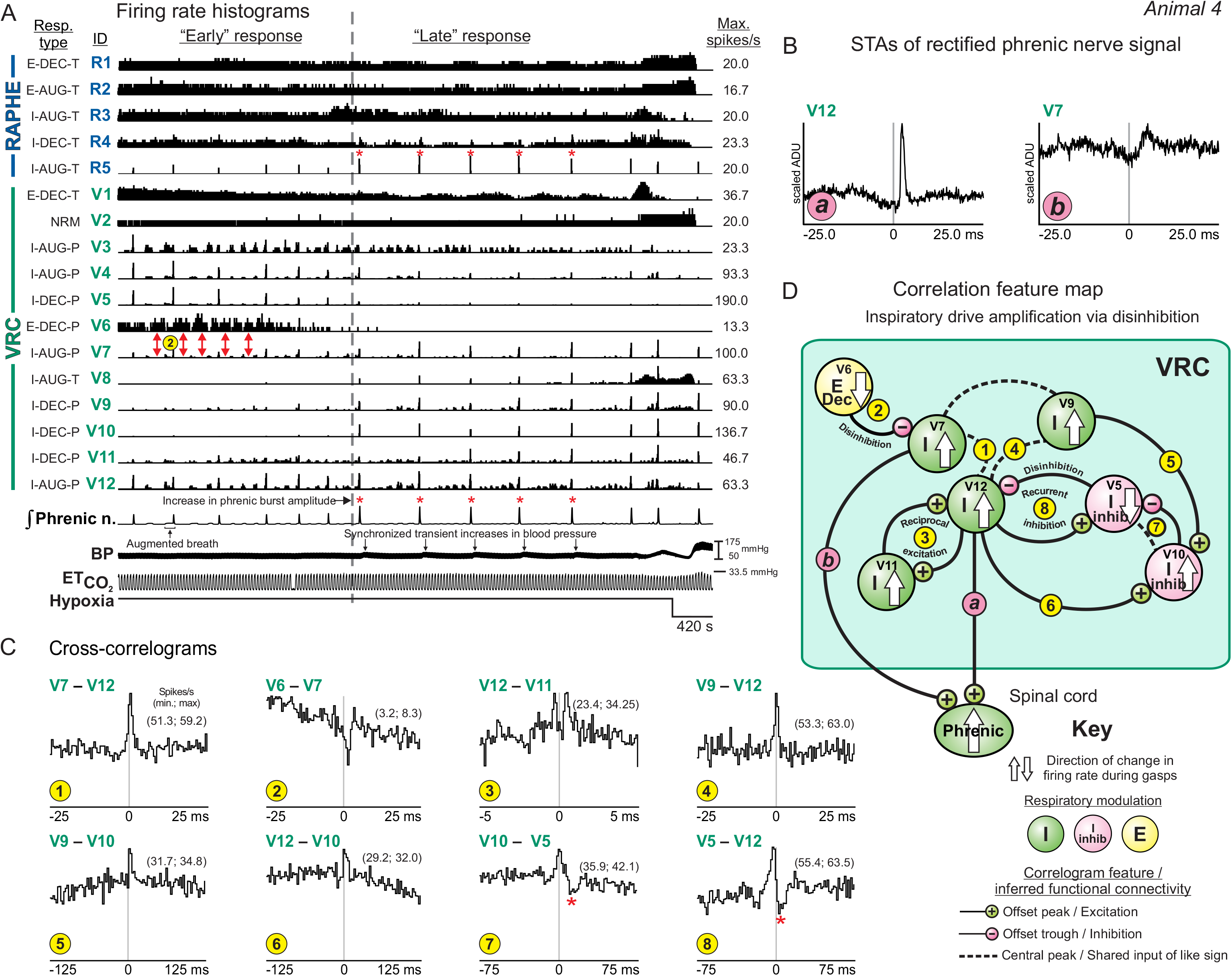
Sequence of phrenic motor discharge patterns and “early” vs. “late” gasp-synchronous bursting neuron groups during ventilation with 5% O_2_. *A*. Firing rate histograms showing detail of firing rates of raphe and VRC neurons during hypoxia-evoked gasp bursts. Note the transition from lower amplitude to higher amplitude bursts of integrated phrenic activity (*red asterisks*) and corresponding changes in burst amplitudes of two subsets of “early” and “late” responding VRC and raphe neurons. Synchronized step-wise increments in systemic arterial blood pressure were associated with each of the “late” response gasp bursts (black arrows). Red double-headed arrows indicate asynchronous firing patterns of cells V6 and V7; functional inhibition of V7 by V6 is inferred by the positive-lag trough feature in cross-correlogram 2 in panel *C*. *B*. STA of rectified phrenic nerve activity triggered by neurons V12 and V7. *C*. Cross-correlograms for the indicated pairs of neurons. DI values for troughs or peaks in each histogram: *1*: 5.8; *2*: 5.77; *3*: 5.00; *4*: 6.96; *5*: 4.38; *6*: 4.1; *7*: 6.71; *8*: 9.42. Number of spikes for each neuron used to calculate cross-correlograms: V10: 86,003; V7: 322,707; V9: 101,717; V11: 103,220; V12: 170,819; V5: 81,986; V6: 43,094. *D*. Correlation feature map for indicated neurons summarizes inferred relationships and indicates mechanisms of interaction and changes in cell firing rate associated with the amplification of inspiratory drive as described in the text.

Coordinated inspiratory neurons V9 and V12 (Fig. 5*C*, 4) each triggered a correlogram with target neuron V10 that featured an offset peak (Fig. 5*C*, 5 and 6), a result consistent with convergent excitation. Neuron V10 was identified as an element of a correlational inspiratory neuron chain; it triggered a correlogram with neuron V5 featuring a positive-lag offset trough relative to a central peak (Fig. 5*C*, 7 *red asterisk*). A similar feature set for pair V5-V12 (Fig. 5*C*, 8) supported the final link in the chain and the hypothesis that it operated via recurrent disinhibition to further amplify the inspiratory drive generated by neuron V12. Inspection of the firing rate histograms of these inspiratory neurons revealed changes consistent with simple interpretations of the short-time scale correlations: as the firing rates of neurons V9 and V10 increased during the late response interval, that of neuron V5 was reduced. The correlation feature map (Fig. 5*D*) summarizes the inferred relationships among these neurons as well as the mechanisms of interaction and directions of change in firing rate associated with the amplification of inspiratory drive as described above.

## Discussion

The results show that acute hypoxia evokes the sequential generation of motor patterns for tachypnea, apneusis and augmented bursts, apnea, and gasping, with corresponding changes in raphe-pontomedullary circuits of the respiratory network. Correlational signatures of neuronal interactions and altered firing rates support the hypothesis that gasp drive is amplified by a disinhibitory inspiratory chain microcircuit and distributed via efference copy to generate concurrent bursts in raphe and pontine neurons, which may be coordinated with step-like increments in arterial blood pressure.

### Tachypnea

Carotid chemoreceptors contribute to the generation of the initial tachypnea with hypoxia, expressed as increases in integrated phrenic nerve amplitude and in respiratory cycle frequency (4). In carotid-deafferented cats, tachypnea generated by central mechanisms is characterized by high frequency and low tidal volumes in an O_2_ concentration-dependent manner (70). In the current study, tachypnea was associated with diverse modulations of firing rate in putative premotor inspiratory neurons. Identified functional connectivity suggested that the tachypneic motor pattern was shaped by tuning of inhibition by expiratory and inspiratory neurons (Fig. 4). The involvement of raphe circuits in the generation of tachypnea has been suggested by prior work (71–73); we observed numerous interactions between raphe and VRC neurons.

### Apneusis

Persistent hypoxia disrupts neuronal metabolism, leading to a general decline in neuronal activity and the transition to prolonged apneustic inspiratory bursts, some ending with fictive augmented breaths or sighs due to cooperative neuronal and glial signaling (9). The apneusis, indicative of delayed inspiratory-to-expiratory phase switching, may be a consequence of diminished glycinergic inhibition of inspiratory neurons by post-inspiratory neurons (74); 5-HT_1A_ receptor agonists can switch this “disturbance” of the respiratory rhythm back to a normal control pattern (75, 76).

### Amplification and distribution of gasp drive

Prior experiments and computational models, particularly work on persistent sodium currents, have documented a functional simplification of the VRC network during the transition to gasp generation (36, 77–81). This is supported by a recent report by Bush and Ramirez, in which many VRC neurons were simultaneously recorded, and neuronal population dynamics were represented in a multidimensional space. Stereotyped VRC “rotational” dynamics were observed during eupnea; these were continuously reconfigured during severe hypoxia, ultimately collapsing into all-or-none ballistic trajectories during gasping, suggesting a dissolution of the normal VRC dynamics as hypoxia progresses to gasping (82). The functional connectivity, altered firing rates, efference copy of gasp drive, and coordinated step increments in blood pressure we report here support a distributed brain stem network model for amplification and broadcasting of inspiratory drive during autoresuscitative gasping that begins with a reduction in expiratory neuron inhibition and an initial loss of inspiratory drive during hypoxic apnea (Fig. 6*; arrows labeled “1”*).

**Figure 6.**
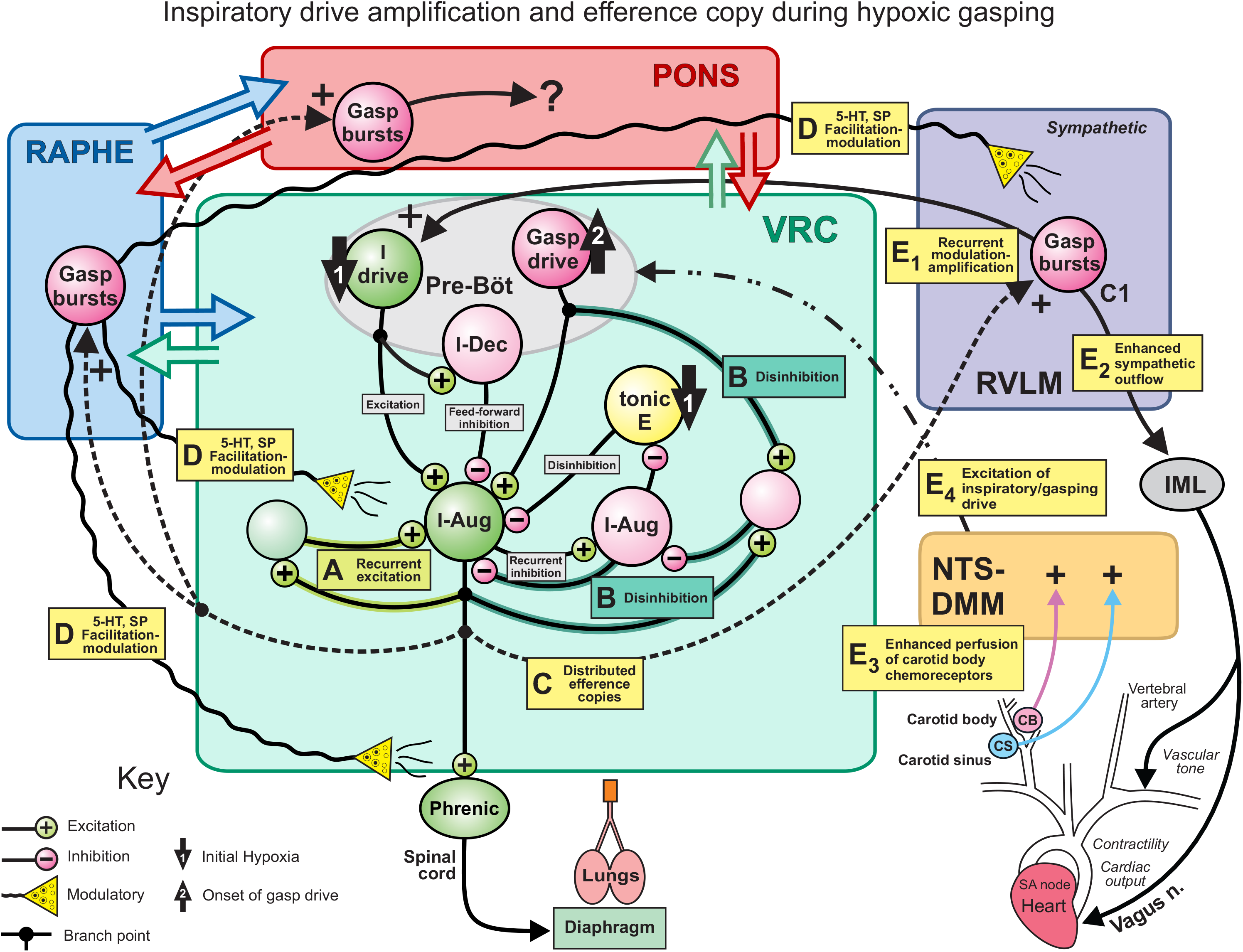
Graphical summary of hypotheses and circuit operations suggested by the results. See text for details. BS, bulbospinal cell CB, carotid bodies CS, carotid sinus IML, intermediolateral nucleus of the spinal cord (origin of preganglionic sympathetic neurons) NTS, nucleus of the solitary tract pre-Böt, pre-Bötzinger complex RVLM, rostral ventrolateral medulla SP, substance P VRC, ventral respiratory column.

The onset of gasp-driver neuron bursting (Fig. 6; *arrow labeled “2”*) excites downstream inspiratory premotor neurons. This onset is due in part to the accumulation of neuromodulators, including 5-HT and substance P released by prior and ongoing raphe activity (40), altered inhibition (83), and factors released by local hypoxia-sensing glia (52, 75, 84, 85). The present results suggest that burst efficacy is further amplified by downstream recurrent excitation (Fig. 6*A*) and disinhibition of inspiration (Fig. 6*B*) within microcircuits of the inspiratory neuron chain. In this context, we identified 24 inhibitory relationships between neurons. Of these, 8 are examples of disinhibition of activity during gasps, i.e., the absence of activity in an inhibiting cell corresponded to gasping activity in the target cell. Seven of the eight neuron pairs included an expiratory VRC cell and a target inspiratory VRC cell. Although the precise source of primary gasp-drive in the cat is not known (86–88), evidence for disinhibitory modulation of inspiratory drive by changes in blood pressure has been reported (50).

Changes in spectral peaks of phrenic nerve signals during the augmented and gasping periods of the response to hypoxia have been reported, such as a shift to higher frequencies during hyperpnea (89, 90), as well as an emergence of power at a lower frequency during gasping (7). We observed changes in time-frequency representations of inspiratory neuron discharge during gasping, as compared to control respiratory cycles (e.g., Fig. 2*F*), consistent with a suppression of recurrent inhibition and the subsequent loss of a distinct signal within the HFO frequency band (i.e., significance is not limited to the 60-110 Hz frequency band).

Efference copies of the inspiratory drive engage raphe, pontine, and rostral ventrolateral medullary (RVLM) pre-sympathetic circuits in a distributed network of coordinated gasp-related bursting (Fig. 6*C; dashed lines*). In turn, burst-synchronous neuromodulator release by raphe neurons further enhances target neurons at multiple brainstem and spinal sites (Fig. 6*D wavy lines*), thereby evoking additional amplifying recurrent circuit loops and the transition from the early low gasp drive state to the later high gasp drive state (e.g., Figs. 4, 5), all of which can potentially contribute to successful autoresuscitation. The gasp drive is amplified by putative recurrent excitation of the pre-Böt I-Driver neurons by C1 RVLM neurons (Fig. 6E1, 9, 37, 91-93) and enhances sympathetic outflows via pre-sympathetic neurons of the RVLM (Fig. 6*E2*). The resulting enhanced coronary function increases perfusion of carotid body chemoreceptors and baroreceptors (12–14) (Fig. 6*E3*), thus increasing the activation of neurons within the nucleus of the solitary tract (NTS) which then relay excitation to the VRC and raphe, respectively (Fig. 6E4, 4). However, the precise functions of the coordinated gasp-synchronous bursting detected in the raphe, as well as the pons, remain unknown.

### Consideration of methods and approach

Parallel spike train recordings from multiple sites within the respiratory brain stem permitted the detection of correlational signatures of neuronal connectivity. Cross-correlation offset peaks and troughs are commonly interpreted as signs of functional excitatory and inhibitory interactions, respectively (94, 95). These features, together with the firing rate modulations associated with the different respiratory motor patterns, suggested several new hypotheses on circuit operations that contribute to the generation of the hypoxia-evoked motor patterns. Advantages and limitations of these approaches have been published (45, 56, 96, 97). Although neurons in the region of the retrotrapezoid nucleus and parafacial respiratory group (RTN-pF) were not monitored in this study, prior work identified evidence for inspiratory drive projections to that region and reciprocal interactions targeting VRC microcircuits, indicating a network architecture with multiple routes appropriate for gain modulation of breathing by central and peripheral chemoreceptors (46, 47, 49, 98).

The cat expresses a wide repertoire of respiratory-related behaviors relevant to human health and is an established *in vivo* model system for studying the mammalian respiratory brain stem (99–102). Decerebration avoids the effects of anesthetics; chemoreceptor-evoked ventilatory responses to hypoxia are similar to those observed in cats that are awake (103).

In the current study, gasping did not change ventilatory efforts because artificial ventilation was used and the cats were neuromuscularly-blocked. To evaluate the short-term consequences of improved gas exchange on the respiratory motor pattern, air was reintroduced into the ventilator and blood pressure was supported to maintain adequate perfusion pressure to the brain and other organ systems. We did not dissociate the extent to which a reduction in blood pressure *per se* contributed to altered network activity and associated changes in blood pressure, either through reduced brain stem perfusion or altered baroreceptor modulation of the respiratory network (42, 50, 104, 105).

### Conclusion

We report that hypoxia-evoked gasps are amplified through a disinhibitory microcircuit within the inspiratory neuron chain and a distributed efference copy mechanism to generate concurrent gasp-synchronous discharges in the raphe-pontomedullary respiratory network. The present results and a large body of prior work support the hypothesis that raphe circuits are engaged during the sequential progression of behaviors in response to hypoxia, from the onset of tachypnea to apnea and autoresuscitative gasping. Resulting large tidal volumes and altered intrathoracic pressure can enhance blood flow and sympathetic activity. Collectively, these processes operate to “bootstrap” cardiovascular function and enhance perfusion of carotid body chemoreceptors and their central actions on the brain’s cardio-respiratory control network, and potentially facilitate resuscitation and recovery during cardiac arrest (1, 11) or seizures (2, 106). The responses of neurons to hypoxic apnea, their functional connections, efference copy of gasp drive, and coordinated step increments in blood pressure support a distributed brain stem network model for enhancing and expanding inspiratory drive during gasping that begins with disinhibition by expiratory neurons and initial decrease in inspiratory drive.

## Acknowledgements

We thank Kim Ruff Carpintier for her excellent surgical skills, and Peter Barnhill, Kathryn Ross, and Andy Ross for their technical support.

## Notes

Kimberly E. Iceman’s current affiliation: Department of Speech Language and Hearing Sciences and Dalton Cardiovascular Center, University of Missouri, Columbia, Missouri.

